# Systematic comparison of ranking aggregation methods for gene lists in experimental results

**DOI:** 10.1101/2022.01.09.475491

**Authors:** Bo Wang, Andy Law, Tim Regan, Nicholas Parkinson, Joby Cole, Clark D. Russell, David H. Dockrell, Michael U. Gutmann, J. Kenneth Baillie

## Abstract

A common experimental output in biomedical science is a list of genes implicated in a given biological process or disease. The results of a group of studies answering the same, or similar, questions can be combined by meta-analysis to find a consensus or a more reliable answer. Ranking aggregation methods can be used to combine gene lists from various sources in meta-analyses. Evaluating a ranking aggregation method on a specific type of dataset before using it is required to support the reliability of the result since the property of a dataset can influence the performance of an algorithm. Evaluation of aggregation methods is usually based on a simulated database especially for the algorithms designed for gene lists because of the lack of a known truth for real data. However, simulated datasets tend to be too small compared to experimental data and neglect key features, including heterogeneity of quality, relevance and the inclusion of unranked lists. In this study, a group of existing methods and their variations which are suitable for meta-analysis of gene lists are compared using simulated and real data. Simulated data was used to explore the performance of the aggregation methods as a function of emulating the common scenarios of real genomics data, with various heterogeneity of quality, noise level, and a mix of unranked and ranked data using 20000 possible entities. In addition to the evaluation with simulated data, a comparison using real genomic data on the SARS-CoV-2 virus, cancer (NSCLC), and bacteria (macrophage apoptosis) was performed. We summarise our evaluation results in terms of a simple flowchart to select a ranking aggregation method for genomics data.

## 1 Introduction

In biology, there are usually many results from different sources for the same or a similar problem. Many results take the form of a list of genes or proteins, especially for screens of genes, transcripts and proteins related to a specific biological process. In almost all cases these gene lists overlap with results from other experiments. Meta-analysis aims to combine the individual gene lists resulting from individual studies to obtain a more reliable answer. Meta-analysis of this kind of data can often be seen as a ranking aggregation and there are many methods for carrying it out [1].

There are various existing methods for ranking aggregation. This study focuses on the unsupervised and rank-based methods since for transcriptomic and genomic level data there are usually no high-quality training data sets with reliable target labels or universally accepted methods for quantification across different data sources [1, 2].

Unranked lists are common in biology. Examples include annotated pathways (e.g. KEGG [3], Reactome [4], Wikipathways [5]), co-expression clusters (e.g. FANTOM5 [6], STRING database [7]) and reports providing a group of entities as the positive result of a study without ranking them. However, unranked lists are not explicitly accommodated by many aggregation methods and are often either excluded from meta-analyses or incorporated in an ad-hoc manner.

All methods investigated in this study can deal with ranked lists as input (reported genes with order information), irrespective of whether each list include all possible entities or not. However, only a few methods among them can accept unranked lists as input. Some approaches like MAIC [8] and Vote-Counting [8] explicitly claim that they can deal with unranked data. Some methods which are designed only for ranked lists can also tackle this type of input with a slight change to the algorithm, such as Borda’s methods [9]. An unranked source can be a special case of ranking with ties (entities with the same ranking) [9] so that methods that can deal with ranking with ties can also accept unranked sources. An example can be RepeatChoice [10], which break ties starting from an input ranking using order information of other sources. A study [9] on ranking with ties modified some methods to adapt them to ties, like Borda’s methods. Borda’s methods have variations using statistics like mean value(MEAN, GEO) or median(MED)[11, 12] and can accept an unranked list by assigning the same ranking to entities within the list, which is intuitively reasonable for unranked sources. In contrast, unranked sources are not explicitly accommodated by some relatively complicated methods although ideas about dealing with absent ranking information especially for ranked top-d partial lists are sometimes proposed. Examples can be Bayesian methods like BiG [13] and BARD [14], with a solid Bayesian theoretical framework and an estimated distribution instead of only a ranking as the result for each algorithm.

The dataset and the problem at hand can influence the applicability of an algorithm a lot. For example, the inclusion of unranked sources can make methods that only accept ranked sources unusable. Noise level and heterogeneity of the noise are also important properties of the data sources and not every method can perform well on very noisy partial ranked data [15]. Optimising a metric which treats all sources equally in the calculation, such as the MEAN method of Borda’s methods, is not suitable when significant noise is included [15]. Moreover, a large number of elements (e.g. around 20000 genes for humans) also makes methods like Cross Entropy Monte Carlo (CEMC) [16] unsuitable because of the computational cost [1]. Considering the features of the genomics datasets, specific evaluation of ranking aggregation methods is thus required to establish their performance.

Ranking aggregation methods usually use simulated datasets for evaluation [1, 14, 15, 17] as there is a lack of large genomics real datasets with definitive answers. In contrast, simulated data provide ground truth values and can also be used to investigate the effect that specific properties of the real data, like the amount of noise or heterogeneity in the source quality, have on the different algorithms.

In many existing studies, including RRA [15], BARC [17], BIG [13], BIRRA [18] and MAIC [8], simulated data was used. Existing research explored datasets with some features of genomics data, including various partial cases to cut lists, noise levels, heterogeneity of source quality [1, 8], and the way of setting top 5% as truth [15, 18]. But some important features for genomics data, including the difference between the top-ranked genes in the truth set and a large number of entities (like over 20,000 genes for humans), have not been systematically explored. To evaluate MAIC [8], 500 entities were used in the simulated data, and the evaluations in [1, 18] used 1000 entities for the simulated data. Assigning a score for each entity subject to noise from some pre-defined distributions is a common way to generate simulated data. The expected mean score of each entity is an arbitrarily defined constant to show the difference in the significance of genes for the simulation. These scores rank genes from top-ranked genes to bottom, or classify genes into truth group and noise group [13, 15, 17].

In previous research about the MAIC algorithm [8], the data set used in the evaluation is generated by ranking *Z* scores sampled from independent normal distributions given a list specified precision (inverse covariance). This data generation model can control the heterogeneity of list quality and average noise level but was in previous research only used to generate small data sets and short lists (500 potential entities) in order to compare the RRA, MAIC, and Vote Counting(VC) method. It has high potential to generate well-simulated data and forms the basis of the new data simulation method proposed in this study.

We first generated realistic synthetic datasets to simulate real features of biological experimental results. We then used these synthetic data, together with real data from selected fields of biology, to systematically evaluate aggregation methods in a range of conditions expected to be encountered in real-world conditions (mixed vs. ranked data, large data set sizes, heterogeneity, noise).

### Contribution

- Viral infection (SARS-CoV-2), cancer (non-small cell lung cancer; NSCLC), and bacterial infection (modulation of macrophage apoptosis) datasets were collected. Each set used in the assessment is extracted from sources either more highly related to the research question of the corresponding meta-analysis or published in a closer time and is considered to be highly reliable.
- A new simulated data generation method is proposed by analysing 3 real datasets in terms of list number, length, quality of sources, heterogeneity of quality, and also the relationship between each significant gene.
- Implementation of investigated ranking aggregation methods and variations of them that are suitable for dealing with genomics data was carried out based on existing source code, enabling them to use the data with the same format easily and providing the ability of using unranked lists as input for some algorithms. They are evaluated systematically on both the real and the simulated datasets.
- The overall evaluation results are condensed in a practical flowchart to select appropriate ranking aggregation methods depending on the ranking information and the heterogeneity in the quality of the data sources, and another version of the flowchart with more information related to the number of included sources to support the selection of methods.

## 2 Results

This section includes the results of experiments on real and simulated data. Only the best-performing methods and some methods with interesting methodology or performance are shown in Figure 2 and Figure 3 for important experiments to avoid clutter in the figures. Full details of the results is provided in the supplement files as plots and tables with all investigated methods included.

### 2.1 Performance measurements

The measurement used to show the accuracy of methods in gene data should be able to weight top-ranked genes more than bottom ones since they are usually more important in biological research. But how important each position is compared with others is not known. Weighting each position can be used when comparing two lists, like average overlap (AO) [19, 20] which provides a score with built-in importance of each position. But the result can be largely influenced by the way a measurement weights each position. To cover the various scenarios in biological research and provide easily understandable results, coverage rates for various cut-offs are selected as the measurements, similar to the measurements used in [1, 13]. Specifically, it is the accuracy value for experiments on simulated data since the ranking for truth is known. Recall-value is used to measure the coverage rate in real data experiments since only unranked truth sets are available. All simulated datasets are generated with 100 repeats for each combination of parameters and 95% confidence intervals are plotted using shading in Figures 2 and 3. To avoid duplicates, the ‘Mix’ version of Borda’s methods and Stuart are only shown in the result of datasets with unranked lists since they perform exactly the same as the corresponding original version in the R Package RobustRankAggreg for ranked lists. The result with top-1000 cutoff is mainly explored, together with a comparison with other smaller cutoffs in supplement files.

### 2.2 Result for datasets with mix of ranked and unranked sources

It can be seen from Figure 2 that MAIC gives the best performance on a large dataset with both ranked and unranked lists like dataset type mixLarge (0m) which emulates SARS-CoV-2 dataset, especially when there is a relatively large heterogeneity of list quality (shown as *D* in simulation). It also gives the best performance for nearly all the cutoffs from top-1 to top-1000 in the evaluation of SARS-CoV-2 datasets. Another one of the best-performing methods for the SARS-CoV-2 dataset is rMixGEO, which reaches the best level when heterogeneity is small and mean noise level is large in the simulated data, like *D* = 0.1 and *M* = 3 whose specific value can be seen in Figure 1. It keeps being one of the best methods when *M* is small for low heterogeneity datasets. As a method using simple statistics, rMixGEO wins against most more complicated methods in this scenario, but it is not as robust as MAIC when heterogeneity is high.

**Fig. 1.**
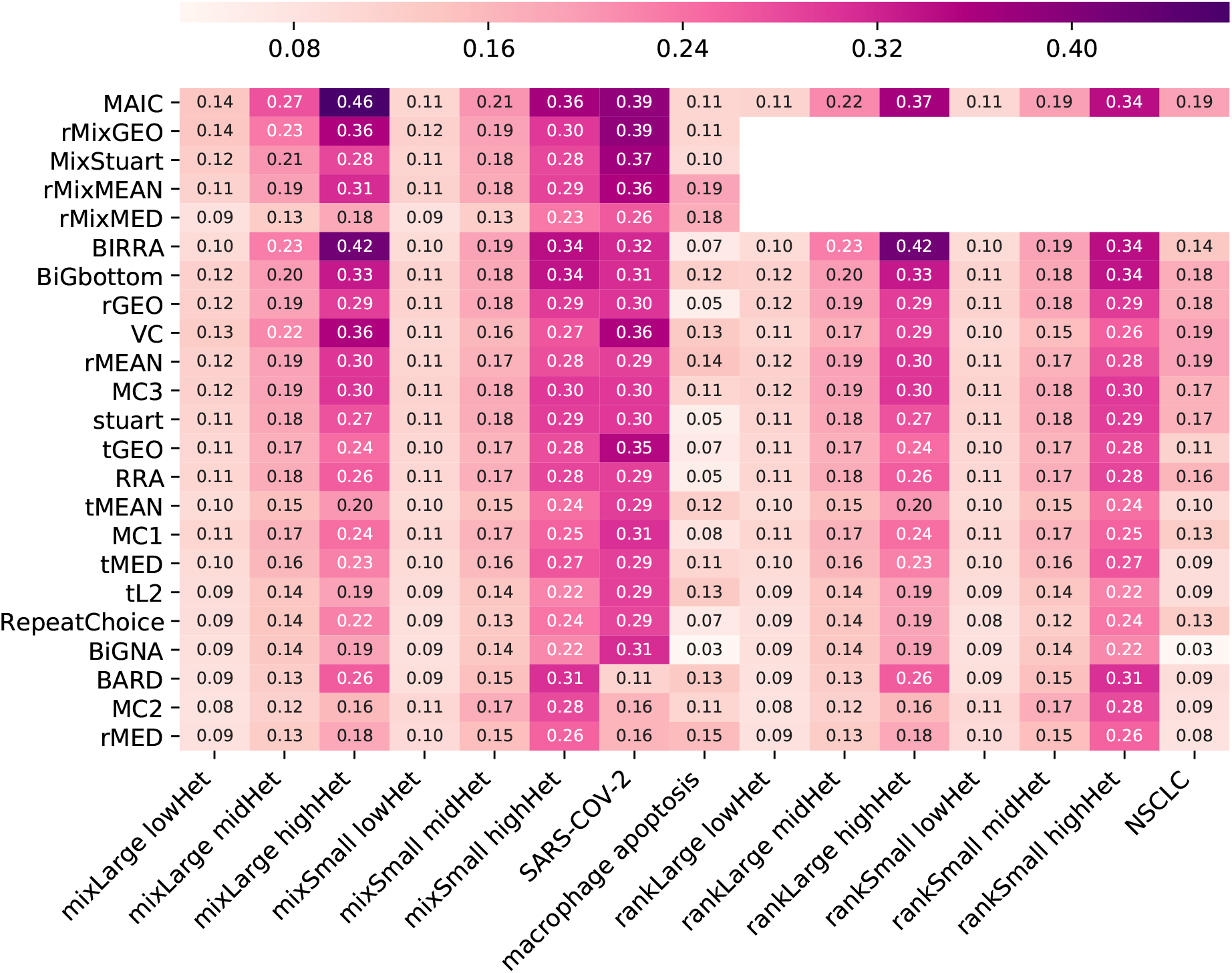
Coverage rates with top-1000 cutoff for important experiments. The name of each investigated method is shown along y-axis. For simulated data, it includes top-1000 accuracy for scenarios with the default mean noise level *M* = 3 and absent gene rate *γ* = 0. The dataset label for simulated data shown along x-axis is the combination of dataset type and heterogeneity. Simulated dataset types are shown as {mixLarge, mixSmall, rankLarge, rankSmall} corresponding to *S* ∈ {0*m*, 1*m*, 0*r*, 1*r*} to show whether unranked lists are included and the number of included lists. The quality heterogeneity is recorded as lowHet, midHet and highHet, corresponding to the small quality heterogeneity *D* = 0.1, the medium one *D* =1 and the large one *D* = 3 separately. For each simulated data setting, the mean value of the results from 100 repeated experiments is plotted. Top-1000 recall for 3 experiments using real datasets(SARS-CoV-2, macrophage apoptosis and NSCLC) are also included.

**Fig. 2.**
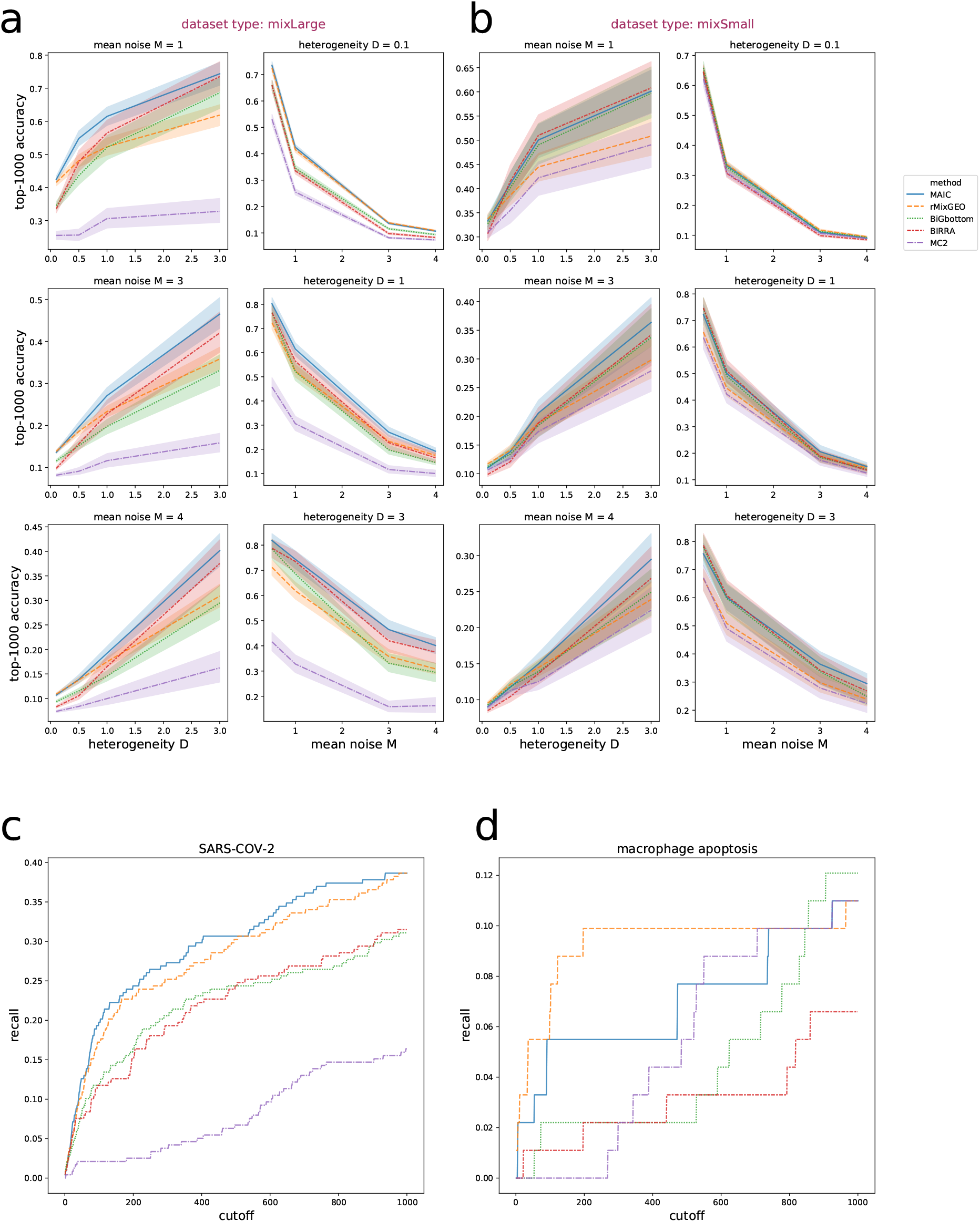
Results for datasets with a mix of ranked and unranked sources. All subplots use the same color and line styles to show investigated methods. (**a**) Simulated data - mixLarge: results for simulated data with various mean noise levels and quality heterogeneity. The simulated dataset type is mixLarge, corresponding to *S* = 0*m* to show that the number of included lists is large. (**b**) Simulated data - mixSmall: results for simulated data with dataset type to be mixSmall, corresponding to *S* = 1*m* to show that the number of included lists is small. For both (**a**) and (**b**), mean of accuracy using top-1000 cutoff and 95% confidence interval are plotted for 100 repeated experiments using lines and shading separately. The default setting of absent gene rate *γ* = 0 is used. (**c**) Real data: results for SARS-CoV-2 dataset. The recall with cutoffs from top-1 to top-1000 is shown. (**d**) Real data: results for macrophage apoptosis dataset. The recall with cutoffs from top-1 to top-1000 is shown.

In terms of a relatively small dataset with a mix of ranked and unranked sources (mixSmall - 1*m* in the simulation), MAIC and rMixGEO follow the same performance pattern as for large datasets. MAIC still reaches the highest performance level whereas rMixGEO experiences a decreasing performance ranking among the investigated methods as the heterogeneity increases. One interesting point is that the performance ranking of BIRRA is influenced a lot by both mean noise and heterogeneity, becoming top-ranked when heterogeneity is high and mean noise is low for small datasets. For each of NSCLC and macrophage apoptosis data, both input lists and the gold standard truth are extracted from less than 10 sources. So the results of NSCLC and macrophage apoptosis data is likely to be quite noisy and less informative than SARS-CoV-2 result. But as the macrophage apoptosis result shows in Figure 2, the results of these real data still identifies that some methods roughly outperform others. rMixGEO and MAIC still show a relatively good performance whereas BIRRA does not perform as well as them. BIRRA and BiGbottom further produce top-ranked results when heterogeneity is high and mean noise is low for small simulated datasets, but they are not as robust to a large mean noise level as MAIC. Plotted in Figure 2, top performed methods including MAIC and rMixGEO are all robust to a change of noise levels under the heterogeneity where they outperform others (all investigated heterogeneity for MAIC and low heterogeneity for rMixGEO) and especially perform well for classic cases (*M* is 3, which is the classic case that emulates real datasets best, see Supplement file 1 - Section 1). They are also robust to various cutoffs and absent gene rates for sources in their corresponding top performed heterogeneity scenarios (all heterogeneity for MAIC and low heterogeneity for rMixGEO), shown as Supplement file 1 - Figure 6, and also the comparison between the result with top-1000 cutoff plotted in Figure 2 and top-100 cutoff plotted in Supplement file1 - Figures 7, 8, and 9.

### 2.3 Result for datasets with only ranked sources

The results for sources with only ranked sources is shown in Figure 3. In this figure, the ranking of BIRRA shows the best performance among investigated methods for large datasets with only ranked sources (rankLarge - dataset type 0r) when heterogeneity *D* reaches 1. The ranking of it tends to be robust to the mean noise level in this scenario. For smaller heterogeneity, rGEO, BiGbottom, MAIC, rMEAN (see Table 1 and Supplement files) and MC3 are top-ranked with similar performance, whereas BiGbottom and MAIC are more robust for high heterogeneity.

**Fig. 3.**
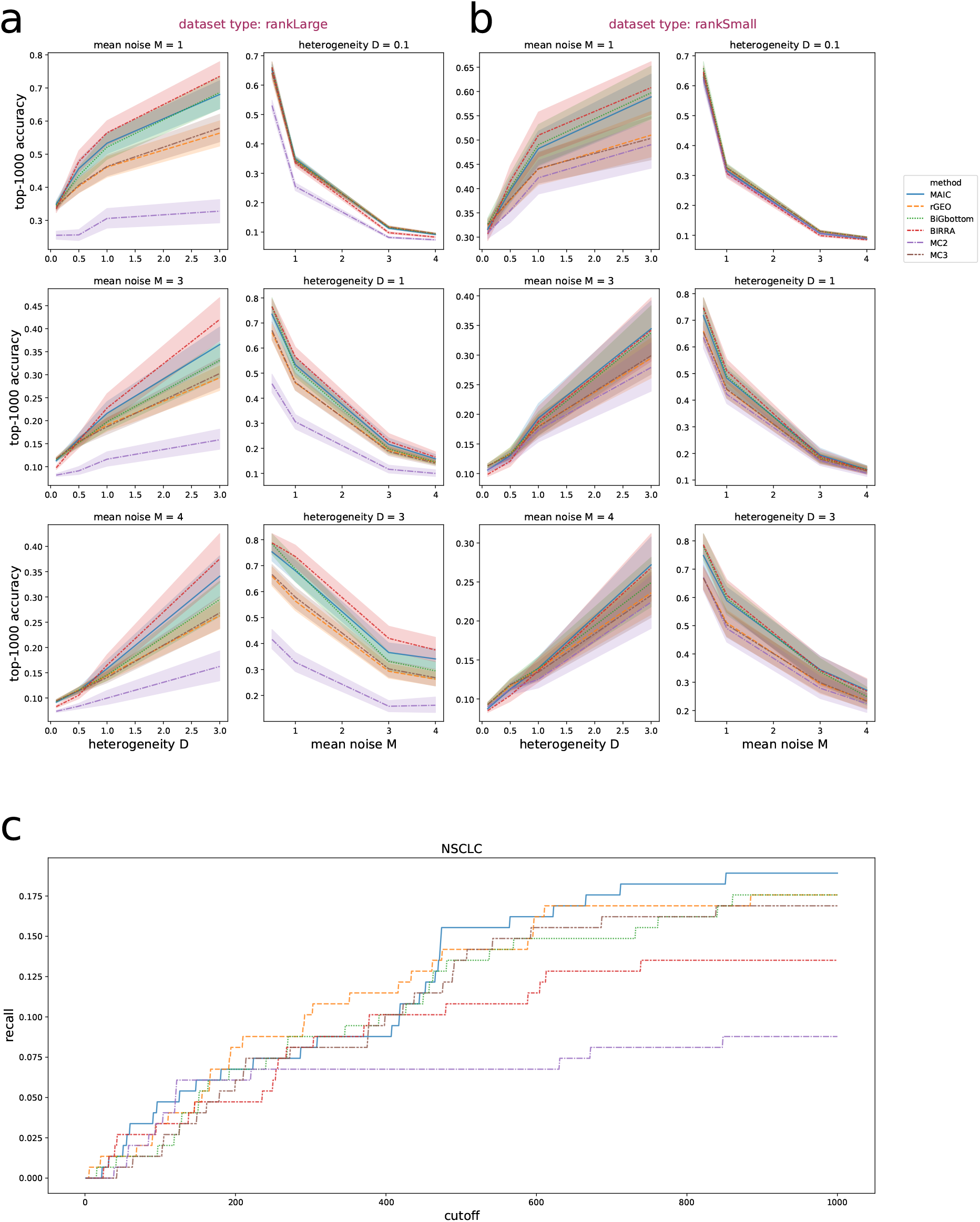
Result for datasets with only ranked sources. All subplots use the same color and line styles to show investigated methods. (**a**) Simulated data - rankLarge: results for simulated datasets with only ranked sources, investigating various mean noise levels and quality heterogeneity. The simulated dataset type is rankLarge, corresponding to *S* = 0*r* to show that the the number of included lists is large. Mean of accuracy using top-1000 cutoff and 95% confidence interval are plotted for 100 repeated experiments using lines and shading separately. The default setting of absent gene rate *γ* = 0 is used. (**b**) Simulated data - rankSmall: Simulated dataset type is rankSmall, corresponding to *S* = 1*r* to show that the number of included lists is small. Any other settings are the same as experiments shown in (**a**). (**c**) Real data: results for NSCLC dataset. The recall with cutoffs from top-1 to top-1000 is shown.

**Table 1.**
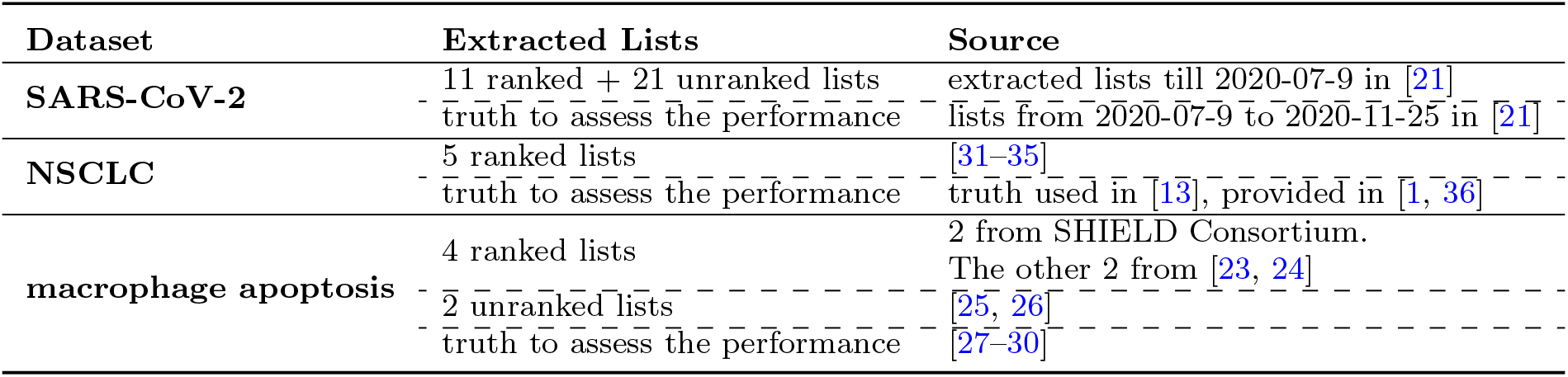
Sources of the collected real data.

Small datasets with only ranked sources (rankSmall - dataset type 1*r*) prefer MAIC, BIRRA, and BiGbottom for high heterogeneity case like *D* is 3, showing the similar best performance for them among investigated methods. Among them, the accuracy of BiGbottom is also one of the highest when heterogeneity is small (*D* equals 0.1), with a similar performance as rGEO and MC3 in this scenario.

Similar to macrophage apoptosis data, NSCLC result also tends to be relatively less informative since the lack of sources for input dataset and truth set. However, MAIC, BIRRA, rGEO, MC3 and BiGbottom all perform relatively well shown as the corresponding plot in Figure 3.

For datasets with a small number of sources, the ranking of BIRRA can be largely influenced by the noise level *M* when heterogeneity *D* is low (around 0.1). But it keeps being one of the best-performing methods when heterogeneity is high. BiGbottom, MAIC, rGEO and MC3 are also robust to the change of mean noise under their corresponding best-performing scenarios of heterogeneity. But MAIC and BIRRA are not as robust in terms of a change of cutoffs for small datasets with high heterogeneity as other best-performing methods. Compared with the results with top-1000 cutoff, they give obvious lower ranked results among investigated methods for accuracy with top-100 cutoff in this scenario, plotted in Supplement file 1 - Section 2.

## 3 Discussion

The difference between the performance of investigated ranking aggregation methods on the proposed simulated datasets and 3 real datasets were compared. The results show that whether to include unranked lists for input data and the heterogeneity of quality for sources can largely influence the performance of the investigated ranking aggregation methods.

The evaluation result in this study can provide some insights on the data selection and method selection for ranking aggregation of genomic problems.

### Data selection

Using a mix of ranked and unranked data instead of giving up unranked sources can lead to a better result, as shown in the simulated data and SARS-CoV-2 results in Figure 1 and the comparison between Figure 2 and Figure 3. When unranked lists are included, the accuracy can grow impressively especially for those top-performing methods, which is shown by the result of MAIC and rMixGEO that accept a mix of ranked and unranked data as input. Their performance improved substantially when unranked lists are included compared to using only ranked sources, reaching the best performance among investigated methods with appropriate heterogeneity. It shows that unranked data has useful information to improve the meta-analysis.

### Method selection

Considering all the evaluation results, a flowchart for selecting methods depending on fundamental properties of the available data is shown in Figure 4. A more detailed version that considers additional information, namely whether the number of sources included is large, is shown in Supplement file 1 - Figure 10. In order to construct the flowcharts, the accuracy with top 1000 cutoff for the result of simulated datasets are firstly considered to select methods in each scenario, followed with a robust checking for the number of sources, various mean noise levels, cutoffs and absent gene rates to only select methods with relatively good robustness for these properties.

**Fig. 4.**
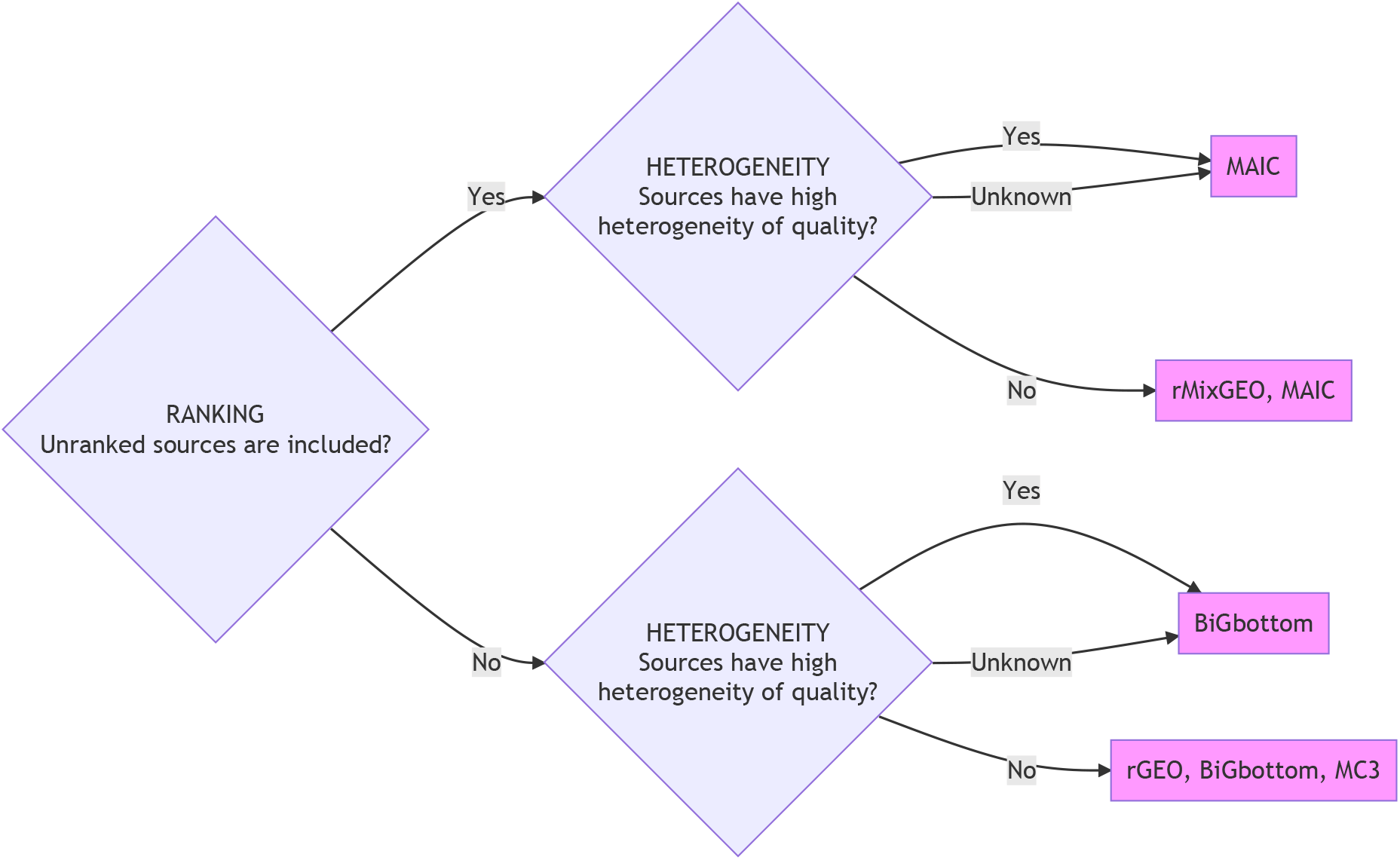
A flowchart for selecting method depending on the ranking information and the heterogeneity of quality for the investigated sources, generated following the evaluation result of this study. Multiple methods within the same block means they perform similar with the best performance under the corresponding scenario.

MAIC and rMixGEO show the best performance among all the methods investigated when using a mix of ranked and unranked data, while MAIC is more robust for the relatively high heterogeneity case. The method rGEO (rMixGEO is a variation of rGEO which can accept unranked lists as input), which is based on simple statistics, is more intuitive and easier to implement and can be selected when the heterogeneity of quality is known to be low, such as a dataset with many sources from the same repeated experiments.

If a dataset only includes ranked data, rGEO, BiGbottom, MAIC, rMEAN, MC3 and BIRRA are preferred for a large dataset with many sources (like dataset type *S* = *r*0). Except for rMEAN, they also show a top-ranked performance for relatively small datasets with only a few sources (like dataset type *S* = *r*1). Among them, the ranking of BiGbottom is more robust for heterogeneity whereas BIRRA performs better among investigated methods for high heterogeneity scenarios.

In terms of the relatively higher robustness to a change in quality heterogeneity for the result of BiGbottom and MAIC, the most likely reason is that they explicitly model and estimate the list quality. Usually, the quality and heterogeneity of quality for input datasets are hard to know, whereas the list type (ranked or unranked) is relatively obvious. So among top-performing methods in the evaluation of this study, methods like BiGbottom and MAIC that parameterize list quality and tend to be robust for various quality heterogeneity are preferred if the heterogeneity is unknown.

General noise level is another property that is usually hard to know in real data. The selected best-performing methods for each dataset type and heterogeneity level shown in Figure 4 are robust to various noise levels under the corresponding scenarios where they give the best performance. Similarly, these selected best-performing methods are also robust to the change of the absent gene rate (*γ*) for sources and the cutoffs for the results (the number of top-ranked genes in the result list used to calculate the accuracy) when calculating the accuracy, shown as Supplement file 1 - Figure 6, with slight fluctuations of their rankings under their best-performed dataset types and heterogeneity. Comparison between top-100 accuracy, which is also plotted in Supplement file 1 - Section 2, and top-1000 accuracy can also suggest the robustness on cutoffs of the results used to evaluate the performance for each method. It can be noticed that the performance ranking of MAIC and BIRRA shows a significant decreasing trend when the cutoff length falls for datasets with only ranked lists and high heterogeneity, especially for small datasets, whereas BiGbottom is more robust for small cutoffs. So although they appear together with BiGbottom as the top-ranked methods for ranked only data with high heterogeneity depending on the results of top-1000 cutoff, BiGbottom is preferred if a study focuses on a small group of top-ranked entities within the result. Considering the robustness of cutoffs, only BiGbottom is selected as the final best-performing method in the Figure 4 where the input data only includes ranked lists with high heterogeneity of quality. But MAIC and BIRRA can also be expected to show the best level of performance when a study focuses on more genes in the result, like top-1000 genes.

In this study, datasets with only order information of some entities are evaluated, including the ranked or unranked list of genes. But methods like MAIC and BiG can also take the classification of sources as additional input information. Classification label can be manually assigned to each source by classifying sources using self-defined criteria like experiment methods or cell types used. So it could be additional information that can be relatively easy to provide. In the future, the influence of classification on ranking aggregation methods and methods for classifying sources will be explored.

## 4 Methods

### 4.1 Real data collection

Three real datasets are collected, corresponding to virus (SARS-CoV-2), bacteria (macrophages apoptosis) and cancer (NSCLC), as shown in Table 1.

The virus data comes from a SARS-CoV-2 related meta-analysis study for prioritisation of host genes implicated in COVID-19 [21]. The dataset used in the evaluation to assess the result is selected from the lists collected by researchers later than the first published version (2020-07-9 to 2020-11-25) and is not used as input for the ranking aggregation algorithms. 238 genes appearing more than once within the top 100 genes of new lists after the published lists (2020-07-9) are extracted as truth. Details of each source for SARS-CoV-2 data can be found in [21].

The cancer dataset consists of genes associated with non-small cell lung cancer (NSCLC) which were collected and used for evaluating ranking aggregation methods by Li et al. (2019, 2018) [1, 13]. 5 lists were extracted from sources that were used in these two papers. The evaluation list used as the truth set in [13] is used to assess the performance of the ranking aggregation methods.

The third dataset concerns genes related to apoptosis in human macrophages during bacterial infection. Two of the lists are provided by the SHIELD Consortium (SHIELD AMR Research Consortium: https://shieldamrresearch.org/) including a ranked list from an infection-associated apoptosis CRISPR screen in human iPSC-derived macrophages and another ranked list of human orthologues of upregulated genes in murine bronchoalveolar lavage alveolar macrophages (BAL AM) 16 hours after infection with Streptococcus pneumoniae [22]. Whereas the other lists for this macrophage apoptosis data set are collected from published papers including [23–30].

The dataset used as the gold standard to assess the performance of the methods is extracted from multiple sources which are either more highly related to the research question of the corresponding meta-analysis or published more recently. It is the same case for all three real datasets. So these sources used as the gold standard are assumed to be relatively reliable and combined as an unranked list for each of the three collected real dataset to assess the performance of the algorithms.

### 4.2 Simulated data and experiment design

#### 4.2.1 Stochastic generative model for the simulated data

In order to better explore the properties of real data, which usually includes a large number of entities, a new stochastic generative model that emulates real data is proposed. It includes 20000 potential entities (human genome scale) in total and incorporates heterogeneity in the lengths of lists. The top 1000 (5%) entities will be considered as signal/significant entities, which corresponds to the number of top-ranked genes being focused on in real research.

For list *L_i_*, entity *E_k_* (k=1…20000), mean noise scale *M*, mean cutting point *m_c_*, heterogeneity of noise *D*, heterogeneity of cutting point *D_c_*, range for each list length [*L, U*], entity significance *μ_k_* for entity *E_k_*, the score *Z_ki_* for entity *E_k_* in list *L_i_* is

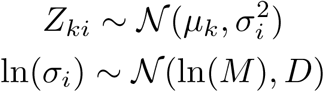

where 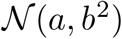 denotes a Gaussian distribution with mean *a* and variance *b*^2^. The number of entities which will be finally included in list *L_i_* is denoted by *N_i_* ∈ [*L, U*] and generated as follows:

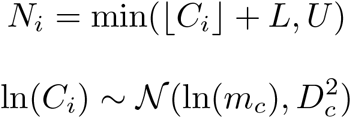

⌊*C_i_*⌋ + *L* is the cutting point for list *L_i_* to only keep the top *N_i_* entities in the list and *N_i_* ∈ [*L, U*]. ln(*C_i_*) is used instead of *C_i_* to make the perturbation on larger scale easier than smaller scale, because the difference on larger scale of length is considered to be less significant. For example, the difference between length 2 and 102 are more significant than the difference between length 19000 and 19100. After the generation, all the potential entities within list *L_i_* are ranked by score *Z_ki_*. Then, *X* entities will be removed randomly from list *L_i_* as controlled by a ratio *γ*,

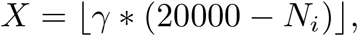

followed by removing the bottomed ranked entities until there are only *N_i_* entities left. Then list list *L_i_* is labeled as “RANKED” or “UNRANKED” to indicate whether the order information for these *N_i_* entities are provided or not. The settings of the hyperparameters of the model, i.e. *L, U, m_c_*, *D_c_, M*, *D*, *μ_k_*, *γ*, the number of the lists and whether each list is ranked or unranked, are discussed in Section 4.2.2 and the Section 1 of Supplement file 1.

In a real study, the length of the reported list is usually a result of two factors. The first one is that bottom-ranked genes are removed and not reported, whereas the second one is that some genes are not included in the study in the first place when the study is not genome-wide. These two situations were both explored by Li et al. (2019) [1] and the new proposed model outlined above. The first case is emulated by removing bottom entities for simulated lists whereas the second case is emulated by removing entities (uniformly) at random.

#### 4.2.2 Parameter settings for new model and comparison with real data

The collected real datasets for SARS-CoV-2, NSCLC, and macrophage apoptosis were explored to set the parameters of the generative model so that it can well emulate the real scenarios. The details about this exploration and parameter settings are included in Section 1 of Supplement file 1. After the exploration of the real data, 4 groups of datasets were generated as shown in Table 2. In the name of dataset types (*S*), “r” means it only includes ranked sources whereas “m” refers to a mix of ranked and unranked sources. In addition, “0” and “1” means large dataset and small dataset respectively. For each of the 4 types of data, various values for *M, D*, and *γ* are used to generate different datasets that cover different scenarios of noise levels, heterogeneity and absent gene rates. By analysing the mean and standard deviation of the accuracy of real sources, *M* ∈ {0.5, 1, 3, 4, 12} and *D* ∈ {0.1, 0.5, 1, 3, 12} were used in the evaluation for each dataset type..

**Table 2.** Different types of simulated data and the corresponding real datasets that they emulate.

| Dataset type ($S$) | Component | Emulated real dataset |
| --- | --- | --- |
| 0r | 11 ranked lists | large dataset: SARS-CoV-2 |
| 0m | 11 ranked lists + 21 unranked lists |  |
| 1r | 5 ranked lists | small datasets: NSCLC, macrophage apoptosis |
| 1m | 5 ranked lists + 5 unranked lists |  |

The different types are distinguished by the length of the list (small "0" or large "1") and whether unranked lists are included ("m") or not ("r").

Shown as Table 3, the properties of real data were explored to set the parameters of the generative model. List number, list length, frequency decay of top-ranked entities, list quality and absent genes of 3 collected real datasets were assessed and the model parameters were set such that the simulated data well emulates them. The details about assessing the real data and the settings of related parameters for the generative model are shown in the Section 1 of Supplement file 1.

**Table 3.** Table for the correspondance between the properties of real data and parameters of generative model.

| Real data | list length boundary | list length distribution | frequency decay of top-ranked entities | list quality | absent genes |
| --- | --- | --- | --- | --- | --- |
| Simulation | $L, U$ | $m_c, D_c$ | $\mu_k$ | $\sigma_i, M, D$ | $\gamma$ |

The parameters were set to emulate properties of the real data.

### 4.3 Selection and implementation of existing methods

As summarised in Table 4, 23 methods have been implemented, based on existing code if accessible. For each method, the table introduces method name, important properties, the new implementation in this study besides enabling them to deal with data in the same format, parameter settings, and if using unranked lists. The implementation language for MAIC, VC, and RepeatChoice is Python whereas it is R for all other investigated methods in Table 4 except for BARD, which has available code implemented in C++. RepeatChoice and VC are implemented based on the algorithms introduced in [10] and [8] respectively, whereas other investigated methods are all implemented based on the available code.

**Table 4.**
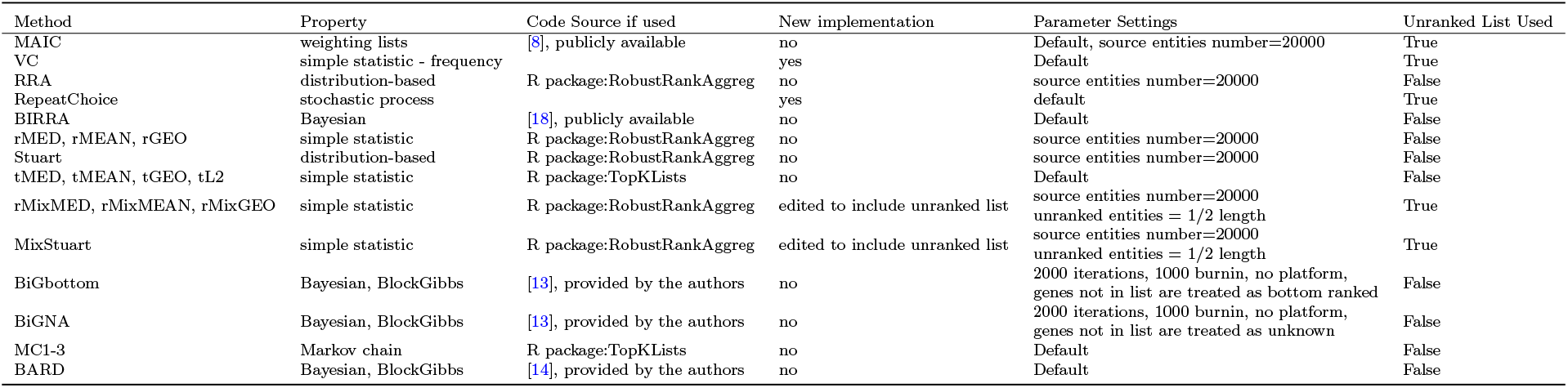
Table for methods investigated in this study.

These methods are selected to include various properties of ranking aggregation methods that are suitable to deal with genomics data. The earliest methods use simple statistics and Borda’s methods are the most popular cases [11, 12]. This type of methods calculate statistics like the mean rank of each entity to get the aggregated result rank. Similar to the research by Li et al. (2019) [1], Borda’s methods are labelled with ‘r’ and ‘t’ to show different implementations from the RobustRankAggreg package and TopKList package. For Borda’s methods and Stuart algorithm from R package - RobustRankAggreg, each of them can easily be edited to enable the input of unranked lists. They are edited by setting genes in unranked lists to have the same ranking, which is half of the list length. The new names for them are rMixMED, rMixMEAN, rMixGEO and MixStuart. These ‘Mix’ versions of Borda’s methods work exactly as the original implementations in the RobustRankAggreg package when dealing with a dataset with ranked sources only.

Bayesian methods are becoming more popular these years [13, 14, 17, 18]. BIRRA [18] initially marks a group of entities as truth and repetitively updates this set. BARD [14] defines a model with an independent parameter for each list to control the probability of position for true entities among noise entities within the corresponding list. In contrast, BiG [13] and BARC [17] use a Thurstone-based model [37], which assumes there is a parameter for each entity and each ranking source is ranked by these parameters subject to noise. Given the solid Bayesian theoretical framework provided, the results can usually be shown in form of an estimated distribution instead of only a single ranking. But it often provides a result largely influenced by priors which are manually defined by the user or specific method. The computational cost is also likely to be large since Monte Carlo methods are usually used to estimate the posterior. BARD and BiG methods are more time consuming than other methods except for Markov chain methods (MC1-3) that can also take days to run for a large dataset. In order to complete each running within or around a reasonable time (48 hours), the same cutting method is used as in [13] and [1]. Before running BARD, BiG (BiGbottom and BiGNA) or MC1-3 on a dataset, genes not ranked within the top 1000 in any list are removed because they are less likely to be ranked within the top 500 in the final result. After this operation, the number of entities in the SARS-CoV-2, NSCLC, and macrophage apoptosis data set became 4584, 2495, and 3267 respectively. The maximal number of iterations in BiG algorithms is set to be 2000 to reduce the running time. Supplement file 1 - Figure 5 shows the comparison between the result of using 5000 iterations and 2000 iterations for BiGbottom and BiGNA. They tend to show the same result so that reducing the number of iterations is not likely to influence the accuracy of the BiG algorithm much and is hence reasonable given the reduction in compute time. Because category information is not investigated in this study, a no-platform version of BiG is applied for both BiGbottom (treating absent genes as bottom-ranked) and BiGNA (treating the order information of absent genes as unknown). Markov chain methods set each unique entity within all input lists as a state and rank entities by the probability of their stationary distributions [38].

In addition to the Bayesian and Markov chain methods, there are other methods that also use the distribution of input rankings or propose a model with distributions like Normal distributions to model the noise which causes the variability within a list, like Stuart [39] and Robust Ranking Aggregation(RRA) [15]. The R Package RobustRankAggreg allows the user to set the number of entities and this parameter in all related implementations is set to be 20000 to emulate the human genome scale, including RRA, Stuart, rMED, rMEAN, rGEO, and the Mix version of Stuart and Borda’s methods.

MAIC also allows to set the number of entities which were again set to 20000. Considering the variation of source quality, some methods try to recognise the difference between lists by weighting them using pre-defined weighting. Pre-defined weighting schemes like in CEMC [16] requires information in addition to the order information, which is usually not available. Methods like MAIC [8] define latent variables to weighting the quality of each source and estimate them without additional information.

## 5 Code availability

The code for simulated data generating and running edited version of algorithms: https://github.com/baillielab/comparison_of_RA_methods/

## 6 Acknowledgments

This work was supported in part by funds from the MRC SHIELD consortium (DHD PI) investigating novel host based antimicrobial responses to antimicrobial resistant bacteria (grant MRNO2995X/1) and Edinburgh Global Research Scholarship.

## 7 Supplementary information

Supplement file 1: The exploration for parameter settings of the stochastic generative model, some result figures of the evaluation and a detailed flowchart for the methods selection.

Supplement file 2: “2_accuracy_M_D_initial_explore.csv” Result table for exploring simulation parameters about *M* and *D*.

Supplement file 3: “3_accuracy-C_plot_D_gamma_average.csv” Result for evaluation on simulated data with various cutoffs of result and absent gene rate *γ*, plotted as figure 6 in Supplement file 1. The mean value of 100 repeated experiments is recorded.

Supplement file 4: “4_top-100 accuracy-M_plot_D_all.csv” Result for evaluation on simulated data for 23 methods and variations. Accuracy with 100 cutoff for the evaluation of datasets including *M* ∈ {1, 3, 4} for *D* ∈ {0.1, 0.5, 1, 3, 12}. Results for 100 repeated experiments are included.

Supplement file 5: “5_top-100 accuracy-D_plot_M_all.csv” Result for evaluation on simulated data for 23 methods and variations. Accuracy with 100 cutoff for the evaluation of datasets including *D* ∈ {0.1, 0.5, 1, 3} for *M* ∈ {0.5, 1, 3, 4, 12}. Results for 100 repeated experiments are included.

Supplement file 6: “6_top-1000 accuracy-M_plot_D_all.csv” Result for evaluation on simulated data for 23 methods and variations. Accuracy with 1000 cutoff for the evaluation of datasets including *M* ∈ {1, 3, 4} for *D* ∈ {0.1, 0.5, 1, 3, 12}. Results for 100 repeated experiments are included.

Supplement file 7: “7_top-1000 accuracy-D_plot_M_all.csv” Result for evaluation on simulated data for 23 methods and variations. Accuracy with 1000 cutoff for the evaluation of datasets including *D* ∈ {0.1, 0.5, 1, 3} for *M* ∈ {0.5, 1, 3, 4, 12}. Results for 100 repeated experiments are included.

Supplement file 8: “8_encoded_lists.zip” Encoded collected real lists and the encoded gold standard used in the evaluation.

## Notes

https://github.com/baillielab/comparison_of_RA_methods

### Competing Interest Statement

The authors have no financial interests to declare. Since our intention is to fairly evaluate a range of methods, it is relevant that J. Kenneth Baillie conceived the MAIC algorithm and Bo Wang, Andy Law and Michael U. Gutmann have worked extensively on development and optimisation of MAIC.

