## Supplement file 1 for "Systematic comparison of ranking aggregation methods for gene lists in experimental results"

#### 1 Parameter settings for the stochastic generative model and comparison with real data

Datasets for SARS-CoV-2, NSCLC, and macrophage apoptosis are used for exploring the features of real data, so that parameters of the stochastic generative model can be selected to better emulate the real data. After the analysis of real data shown as below, parameters are selected for generating simulated data, shown in Table 1 and Table 2.

<table>
<tr><th>list group name</th><th>number of lists</th><th>$\ln(m_c)$</th><th>$D_c$</th><th>M</th><th>D</th><th>$\gamma$</th><th>$\mu_k$, k=1,...,1000</th><th>L</th><th>U</th></tr>
<tr><td>$D1_{ranked}$</td><td>11 ranked lists</td><td>5.3597</td><td>1.8055</td><td rowspan="4">{0.5, 1, 3, 4, 12}</td><td rowspan="4">{0.1, 0.5, 1,3, 12}</td><td rowspan="4">{0, 0.2, 0.5} when M=3, 0 for others.</td><td rowspan="4">$\mu_k \propto a * \exp(b * k) + c$ a=0.1405 b=-0.0099 c=0.0625, $\mu_1 = 2$</td><td rowspan="4">2</td><td rowspan="4">20000</td></tr>
<tr><td>$D1_{unranked}$</td><td>21 unranked lists</td><td>3.4365</td><td>1.6937</td></tr>
<tr><td>$D2_{ranked}$</td><td>5 ranked lists</td><td>7.4141</td><td>1.8634</td></tr>
<tr><td>$D2_{unranked}$</td><td>5 unranked lists</td><td>2.9905</td><td>1.3390</td></tr>
</table>

Table 1: Table for parameter used in simulated data generation.

| Dataset type (S) | Component |
|---|---|
| 0r | $D1_{ranked}$ |
| 0m | $D1_{ranked} + D1_{unranked}$ |
| 1r | $D2_{ranked}$ |
| 1m | $D2_{ranked} + D2_{unranked}$ |

Table 2: Table for the component of each data set used in simulated data generation. For example, a dataset of m0 type includes a group of $D1_{ranked}$ lists and a group of $D1_{unranked}$ lists.

**List number**: The number of lists used in the SARS-CoV-2 study is 32, which includes 21 unranked lists and 11 ranked lists. The small data set used for NSCLC has 5 ranked lists whereas 4 ranked lists are collected for macrophages apoptosis with another 2 unranked lists. Because some unranked sources are used for extracting the gold standard for NSCLC and macrophage apoptosis data, the total number of unranked sources which can be used as input should be higher if collecting data in a similar way, although these 2 datasets are not all sources from a thorough review for related studies. For the simulated dataset, 4 scenarios are analysed. The first two are formal biological meta-analysis analysis cases emulating the collected SARS-CoV-2 data, with 11 ranked lists with or without 21 unranked lists. The other two cases emulate the small collected dataset with 5 ranked lists, and with or without 5 unranked lists.

**List length**: The length of each list can go from 1 to 20000 (human genome scale) theoretically. But studies that included a single gene were excluded in the SARS-CoV-2 study. Also, methods like MAIC are designed for accepting lists with at least 2 entities. So we set a boundary $L = 2$ as the least number of entities in a list. Besides, considering the human genome scale, 20000 is assigned to $U$ as the upper bound to show all generated lists longer than $U$ as a human genome-wide source with 20000 genes.

**Length distribution**: For each list in the study of SARS-CoV-2 with length denoted by $h$, the mean and standard deviation for $\ln(h-2)$ are calculated. The mean for ranked lists is 5.3597 and the standard deviation is 1.8055. So $\ln(m_c)$ and $D_c$ are set to be 5.3597 and 1.8055 separately. The calculated $\ln(m_c)$ and $D_c$ for unranked data lists are 3.4365 and 1.6937, which are the corresponding mean and standard deviation for $\ln(h-2)$. The ranked lists and unranked lists are analysed separately because unranked lists tend to be shorter. Because of the lack of lists for NSCLC and macrophage apoptosis data, especially for the unranked lists in terms of NSCLC data, they are both considered as small data set and analysed together. The histogram and mean value of $\ln(h-2)$ for real data are shown in Figure 1, whereas selected parameters for $\ln(m_c)$ and $D_c$ are shown in Table 1.

**Entity significance** $\mu_k$: In the simulated data generation model, 0 is used as the $\mu_k$ for all noise entities. But the relationship between $\mu_k$ for signal entities needs to be explored.

The frequency of each gene that appears in the top 1000 of all sources was calculated independently for 3 real datasets. For SARS-CoV-2 data, 986 out of 11470 entities show a frequency larger than 0.05 and are considered as signal entities whereas the others are considered as noise. An exponential curve $y_k = a * \exp(b * k) + c$ is fitted for the signal entities ranked by the frequency, with rank $k$ and frequency $y_k$. The $y_k = a * \exp(b * k) + c$ value with these fitted parameters can be used to estimate the significance of an entity($\mu_k$). The same analysis is carried out for NSCLC and macrophage apoptosis data, shown in Figure 2. As the bottom right of the figure shows, the curves for 3 real data sets are scaled to set the first value to be 2 since only the ratio between different positions is investigated here. These curves are similar, all falling from quickly to slower and tending to be flat reaching the 1000th position. So an assumption is made that all these data have a close pattern of significance falling along with the signal genes. The scaled value $y_k$ of the curve for SARS-CoV-2, which is fitted using the most number of lists, is used to set the $\mu_k$ value, which goes from for $\mu_1$ to for $\mu_{1000}$. The fitted curves for the simulated data with $M = 3$ and $D \in \{1, 0.5, 0.1\}$ (explored to simulate appropriate mean noise and variance for simulated data described in the following part) are compared with the curves for 3 real data sets after scaling.

**List quality** $\sigma_i$: Quality of lists in real data was compared to simulated data with various $M$ and $D$, to explore the appropriate settings for parameters $M$ and $D$ which were shown in Table 1. The genes within the corresponding

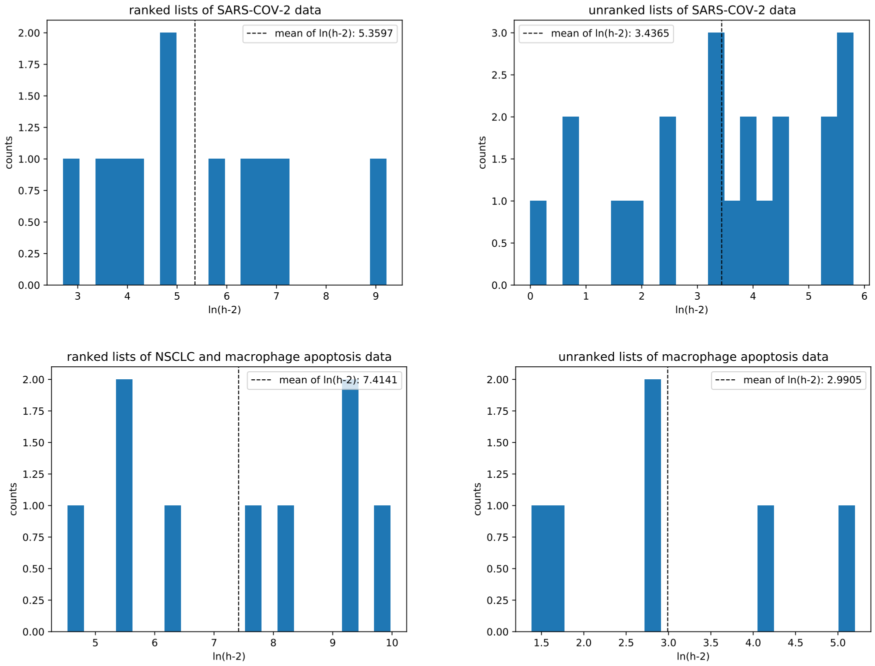


Figure 1: The histogram of $\ln(h-2)$ for SARS-CoV-2 data, and the combination of NSCLC and macrophage apoptosis data, shown as counts and mean value. Figures above are SARS-CoV-2 data. The left bottom one is the analysis of the combination of NSCLC and macrophage data, whereas the right bottom one is for all collected unranked data for macrophage apoptosis, including the constituent of extracted truth.

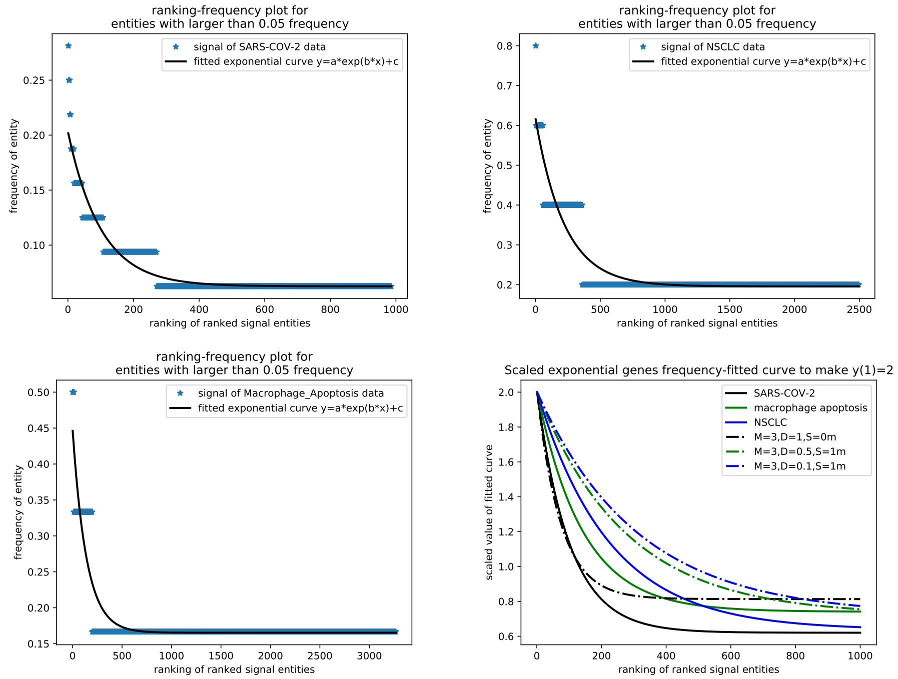


Figure 2: The left and top 3 figures show the frequency for each entity appearing in top 1000 of each list from each real data set except for the pre-selected gold standard list. Entities with frequency larger than 0.05 are considered as signals and ranked by the corresponding frequency. A curve is fitted for each dataset. The right bottom figure shows the plotting for these scaled curves with top ranked entity having value 2, together with the analysis for selected simulated data.

| Dataset | Length of gold standard set | Accuracy | | Upper bound estimation | Lower bound estimation |
|---|---|---|---|---|---|
| | | mean | standard deviation | mean accuracy | mean accuracy |
| SARS-CoV-2 | 238 | 0.1053 | 0.1222 | 0.2993 | 0.0117 |
| NSCLC | 148 | 0.0283 | 0.0263 | 0.3019 | 0.0074 |
| macrophage apoptosis | 91 | 0.0512 | 0.0951 | 0.2124 | 0.0045 |

Table 3: Table for the calculated accuracy for NSCLC, macrophage apoptosis, and SARS-CoV-2 datasets independently. Upper bound and lower bound estimation are also included.

extracted unranked gene set used to assess the performance of algorithms are considered as truth, with $n_T$ genes. In order to assess the quality of each list, the precision($\frac{TruePositive}{PredictedPositive}$) for each list is calculated, which is the ratio for the number of true genes appearing in top $n_T$ genes of a ranked list with more than $n_T$ genes. For an unranked list with length $h$ larger than $n_T$, $n_T$ genes are randomly selected from the list as the predicted positive since there is no order information provided between genes within an unranked list. For a list with $h$ genes less than $n_T$, the precision value is the ratio of true genes appearing in the list. The quality of lists in real data was explored, shown as Table 3. This precision value shows the quality of each list, which is related to the $\sigma_i$ in the simulated data generation model. The average of these precision values is 0.0283, 0.0512 and 0.1053 for NSCLC, macrophage apoptosis, and SARS-CoV-2 respectively. The standard deviations (std) for each of them are 0.0263, 0.0951, and 0.1222. The number of gold standards set $N_T$ for NSCLC, macrophage apoptosis, and SARS-CoV-2 are 148, 91 and 238 separately. Because the size of the truth set may influence the accuracy, these numbers are analysed separately. The unranked lists used as the true positive for these real data are extracted from multiple sources as estimations for the corresponding truth. This estimation can influence the precision values calculated a lot if it is far from the truth. An estimation of the upper bound under various true positive sets with the same length ($N_T$) can be calculated by using the most common positive reported genes as the truth, which are 0.3019, 0.2124, and 0.2993 separately as shown in Table 3. The expected precision value for a non-information random selected list from 20000 genes are $N_T/20000 = 0.0074, 0.0045, 0.0117$ as shown in Table 3, which can be an estimation for the lower bound of precision values.

To explore a suitable noise level $M$ and heterogeneity $D$ used in the simulated data generation model, experiments on various $M$ and $D$ were made to compare with the real data, with other parameters set as Table 1 for a large dataset $m0$ (11 ranked + 21 unranked) and a small dataset $m1$ (5 ranked + 5 unranked) as shown in Table 2. $\gamma = 0$ is used here. $M \in \{0.5, 1, 2, 3, 4, 12\}$ is explored when $D \in \{0.1, 0.5, 1, 3, 12\}$. The no-heterogeneity scenario that $M$ is constant is also explored, shown as $D = 0$. These $M$ value are selected to show various signal noise ratio: $\frac{\mu_k}{\sigma_i}$. Considering the expectation $\mathbb{E}[\ln(\sigma_i)] = \ln(M)$, $\frac{\mu_k}{M}$ is used to show the estimated scale for signal-to-noise ratio($SNR$). Since $\mu_k \in [0.6202, 2]$ when k=1,...1000, 1 is selected for $\mu_k$ to calculate this figure. $SNR \in \{2, 1, 0.5, 0.333, 0.25\}$. Experiments are repeated 10 times and the average of results are calculated for both mean accuracy and standard deviation of accuracy,

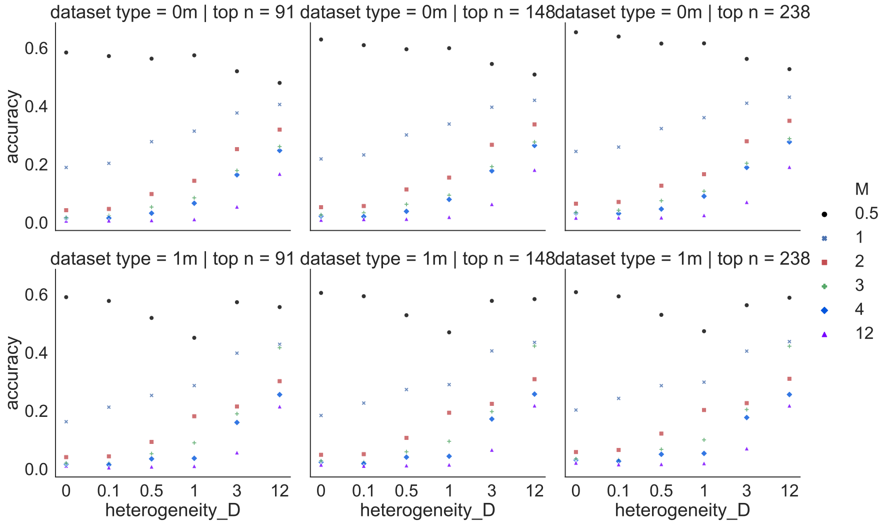


Figure 3: The figure shows the plot for mean accuracy calculated under $M \in \{0.5, 1, 2, 3, 4, 12\}$ and $D \in \{0, 0.1, 0.5, 1, 3, 12\}$. $D = 0$ means $M$ is constant. Each result is an average of 10 repeat experiments.

shown as Figure 3 and Figure 4. The details are shown in Supplement file "2_accuracy_M_D_initial_explore.csv" as a table. The result shows M=3, D=1 best emulates the SARS-CoV-2 data by generating lists with a mean top-238 accuracy of 0.1073 and standard deviation of 0.1858 for the large dataset. For the top-148 accuracy, the mean and std value for NSCLC are 0.0283 and 0.0263. Simulated data with M=3 and D=0.1 has similar values with 0.0245 mean and 0.0275 std value for the small dataset. In terms of the top 91-accuracy, the mean and std value for macrophage apoptosis are 0.0512 and 0.0951. Data with M=3 and D=0.5 has similar values with 0.0525 mean and 0.0694 std value for the small dataset. So M=3 will be treated as a classic important setting when exploring other parameters.

Estimated lower bounds and upper bounds are also compared to explore parameters for emulating a large enough range of various real cases. The smallest mean values for accuracy are around 0.0106, 0.005, and 0.017 for M=12, D=0.1 with the corresponding accuracy for top-$N_T$ cutoff, which are close to the non-information list value (the lower bound estimation) mentioned above and low enough for emulating the lists in real data. It goes to around 0.6 when $M = 0.5$, which is far higher than the upper bound estimation for 3 real datasets and larger than any collected real list accuracy, which tends to be high enough to emulate the real data. In terms of the range of the heterogeneity, the std of accuracy goes from close to no heterogeneity ($D = 0$) when $D = 0.1$ to around 0.4 when D is larger, which is times higher than any real calculated std and

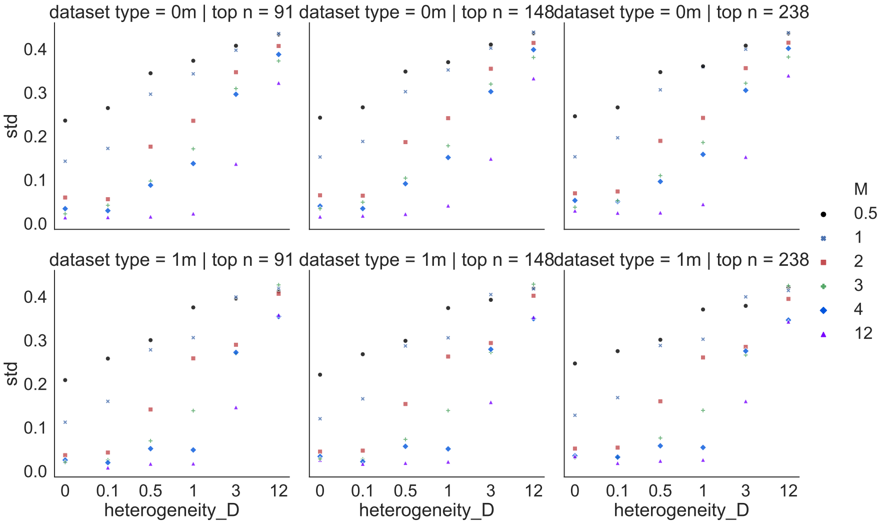


Figure 4: The figure shows the plot for standard deviation of accuracy, calculated under $M \in \{0.5, 1, 2, 3, 4, 12\}$ and $D \in \{0, 0.1, 0.5, 1, 3, 12\}$.$D = 0$ means $M$ is constant. Each result is an average of 10 repeat experiments.

seems high enough in the real case.

Considering the analysis above, parameters settings for $M$ and $D$ used in the evaluation are selected. Various $M$ and $D$ will be used to explore themselves besides $M = 3$ and $D \in \{0.1, 0.5, 1\}$. $M \in \{0.5, 1, 3, 4, 12\}$ and $D \in \{0.1, 0.5, 1, 3, 12\}$ are used to explore various average quality and the heterogeneity of list quality. For heterogeneity $D$, it is the variance of $\ln(\sigma_i)$ with expectation $\mathbb{E}[\ln(\sigma_i)] = \ln(M)$ in the data generation model. When $M = 3$, $(ln(M))^2 = 1.2$. The corresponding signal-to-noise ratio $SNR = \frac{(\ln(M))^2}{D} \in \{12, 2.4, 1.2, 0.4, 0.1\}$ for $D \in \{0.1, 0.5, 1, 3, 12\}$. It can also support that the settings for $M$ and $D$ can generate datasets to cover cases of various mean noise and heterogeneity including common cases in real data for a large reasonable range of $SNR$ for both $\frac{\mu_k}{M}$ and $\frac{(\ln(M))^2}{D}$.

**Absent genes** $\gamma$: For the absence of genes in a real study, it is usually hard to judge the cases between bottom omitted or randomly excluded because the genes included in a study are usually more or less selected using prior knowledge. A simple way to emulate the combination of various scenarios for the exclusion of genes is used by randomly removing 20% and 50% of genes to be removed before cutting the bottom genes for $M = 3$, $D \in \{0.1, 0.5, 1, 3, 12\}$ in terms of 4 types of datasets in Table 2.

### 2 Supplement result figures

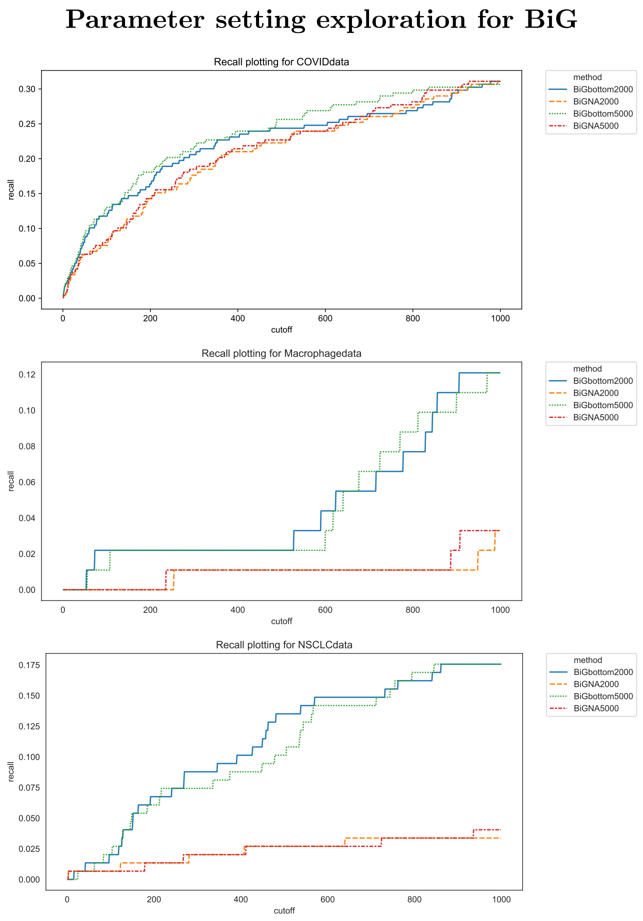


Figure 5: Parameter setting exploration for BiGbottom and BiGNA. The recall for top 1 to 1000 genes in the result list of 3 real datasets for BiGbottom and BiGNA, running with 2000 iterations and 5000 iterations.

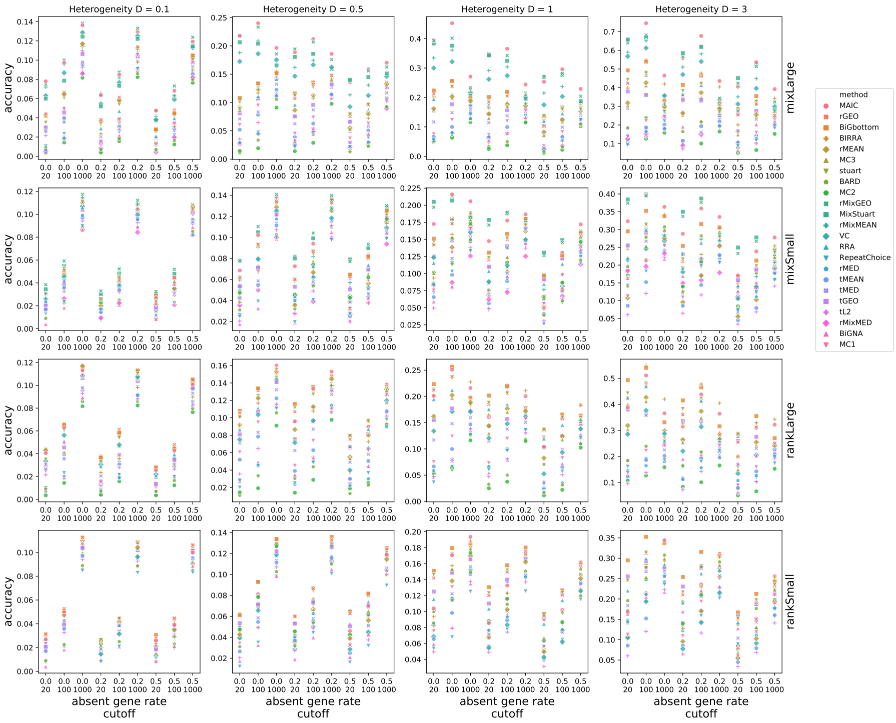


Figure 6: Result for simulated datasets with various absent gene rates and cutoffs. Average value of 100 replicates for each setting is plotted. The accuracy for top 20, 100, and 1000 genes in the result list of simulated data sets are plotted with the change of gene absent rates ($\gamma \in \{0, 0.2, 0.5\}$) when the quality heterogeneity goes from really low to high ($D \in \{0.1, 0.5, 1, 3\}$) and mean noise level emulates the default classic scenario ($M = 3$). Simulated dataset types are shown as {mixLarge, mixSmall, rankLarge, rankSmall} corresponding to $S \in \{0m, 1m, 0r, 1r\}$ to show whether unranked lists are included and the number of included lists.

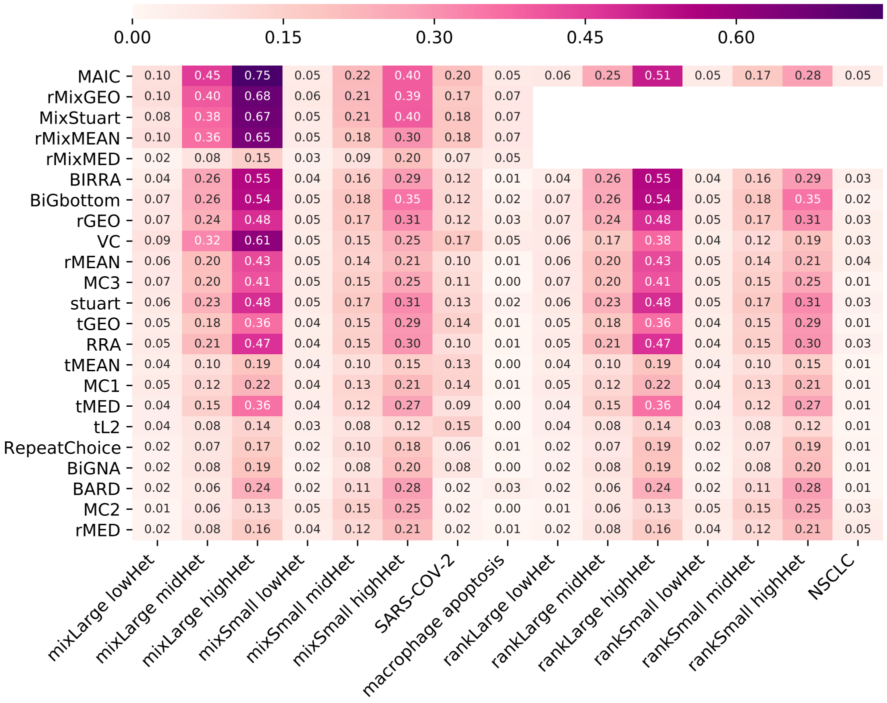


Figure 7: Coverage rates with top-100 cutoff for important experiments. The name of each investigated method is shown along y-axis. For simulated data, it shows top-100 accuracy for scenarios with the default mean noise level $M = 3$ and absent gene rate $\gamma = 0$. The dataset label for simulated data shown along x-axis is the combination of dataset type and heterogeneity. Simulated dataset types are shown as {mixLarge, mixSmall, rankLarge, rankSmall} corresponding to $S \in \{0m, 1m, 0r, 1r\}$ to show whether unranked lists are included and the number of included lists. The quality heterogeneity is recorded as lowHet, midHet and highHet, corresponding to the small quality heterogeneity $D = 0.1$, the medium one $D = 1$ and the large one $D = 3$ separately. For each simulated data setting, the mean value of the results from 100 repeated experiments is plotted. Top-100 recall for 3 experiments using real datasets(SARS-CoV-2, macrophage apoptosis and NSCLC) are also included.

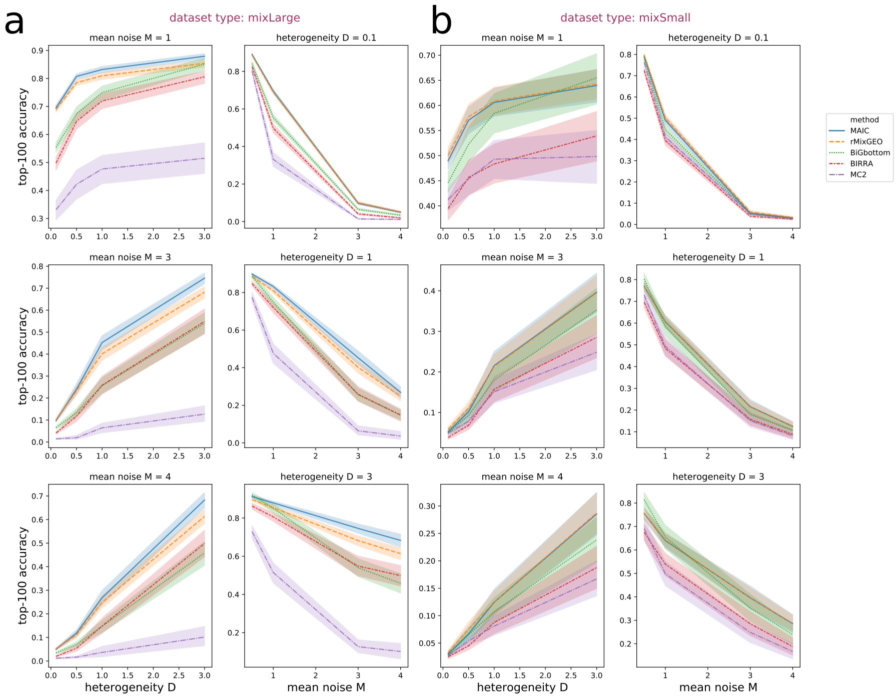


Figure 8: Results for a mix of ranked and unranked sources for simulated data with various mean noise level and quality heterogeneity. Mean of accuracy using top-100 cutoff and 95% confidence interval are plotted for 100 repeated experiments using lines and shadows seperately. The default setting of absent gene rate $\gamma = 0$ is used. (**a**) mixLarge: Simulated dataset types is mixLarge, corresponding to $S = 0m$ to show both ranked and unranked lists are included and the number of included lists is large for each dataset. (**b**) mixSmall: Simulated dataset types is mixSmall, corresponding to $S = 1m$ to show both ranked and unranked lists are included and the number of included lists is small for each dataset.

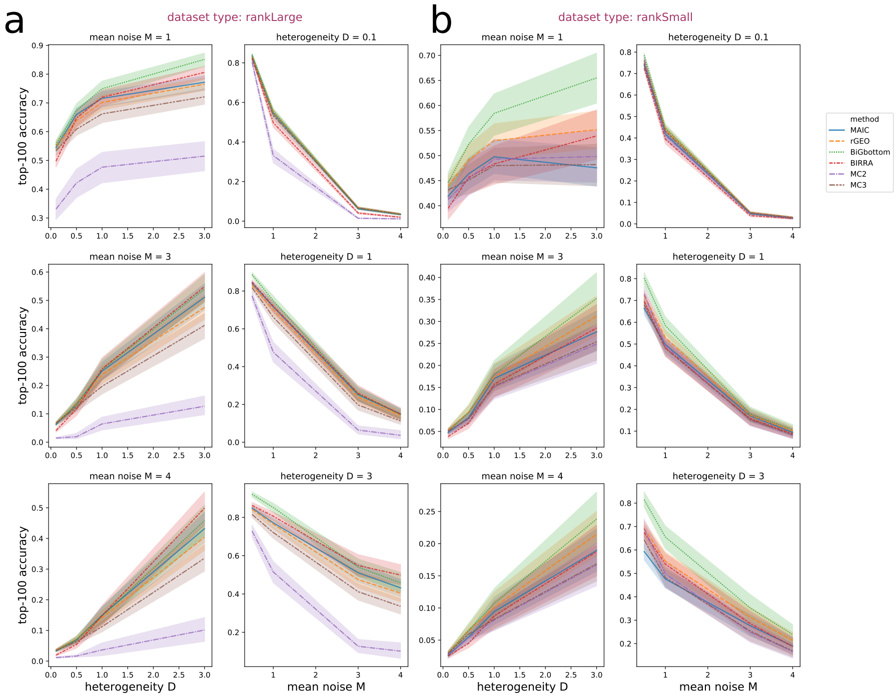


Figure 9: Results for simulated datasets with only ranked sources with various mean noise level and quality heterogeneity. Mean of accuracy using top-100 cutoff and 95% confidence interval are plotted for 100 repeated experiments using lines and shadows seperately. The default setting of absent gene rate $\gamma = 0$ is used. (**a**) rankLarge: Simulated dataset types is rankLarge, corresponding to $S = 0r$ to show that only ranked lists are included and the number of included lists is large for each dataset. (**b**) rankSmall: Simulated dataset types is rankSmall, corresponding to $S = 1r$ to show that only ranked lists are included and the number of included lists is small for each dataset.

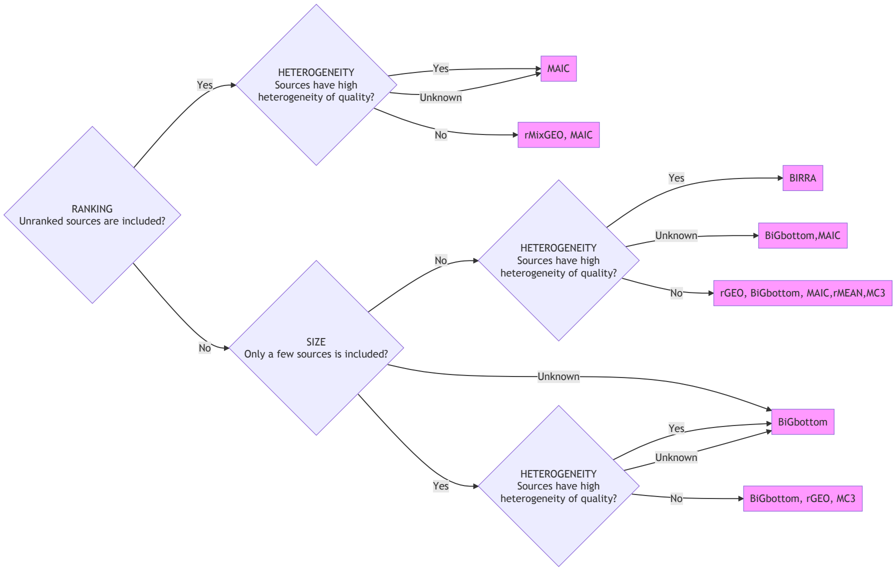


Figure 10: A flowchart for selecting method depending on the ranking information, size, and heterogeneity of quality for investigated dataset, generated following the evaluation result of this study. Multiple methods within the same block means they perform similar with the best performance under the corresponding scenario.
